# Analytic Bounds on GAMLSS Model Variability of Normative White Matter Brain Charts

**DOI:** 10.64898/2026.01.28.702453

**Authors:** Michael E. Kim, Gaurav Rudravaram, Adam Saunders, Chenyu Gao, Karthik Ramadass, Nancy R. Newlin, Praitayini Kanakaraj, Sam Bogdanov, Derek Archer, Timothy J. Hohman, Angela L. Jefferson, Victoria L. Morgan, Alexandra Roche, Dario J. Englot, Susan M. Resnick, Lori L. Beason Held, Murat Bilgel, Laurie E. Cutting, Laura A. Barquero, Micah A. D’Archangel, Tin Q. Nguyen, Kathryn L. Humphreys, Yanbin Niu, Sophia Vinci-Booher, Carissa J. Cascio, Kimberly R. Pechman, Niranjana Shashikumar, The HABS-HD Study Team, Alzheimer’s Disease Neuroimaging Initiative, The BIOCARD Study Team, Zhiyuan Li, Simon N. Vandekar, Panpan Zhang, John C. Gore, Yihao Liu, Lianrui Zuo, Kurt G. Schilling, Daniel C. Moyer, Bennett A. Landman

## Abstract

Brain charts, or normative models of quantitative neuroimaging measures, can identify trajectories of brain development and abnormalities in groups and individuals by leveraging large populations. Recent work has extended these brain charts to model microstructural and macrostructural features of white matter. Assessments of variance for these brain charts are necessary to determine whether the models being used for these data are stable. We implement an analytic approach to characterize variability of the parameters in previously released brain charts created using the generalized additive models for location, scale, and shape (GAMLSS) framework. Additionally, we empirically validate the accuracy of each analytic model through a comparison to a bootstrapping approach from 0.2 to 90 years of age. We find that across all models, the analytic coefficient of variation (COV) remains below 5% for ages greater than 0.25 years, with the maximum empirical observed COV reaching 7% at 0.2 years of age. Further, the empirical assessment shows high agreement with the analytic assessment, with COV estimates averaged across the lifespan for all models having a Pearson correlation coefficient of 0.776 and a mean difference of 4 × 10^−4^. Both methods exhibit volume and surface area as the features with the largest average COV for the majority of tracts. However, the analytic assessment yields axial diffusivity as the feature most frequently having the smallest COV, whereas the corresponding feature for the empirical assessment is average length. These results suggest that the analytic approach overestimates model stability for WM brain charts when the COV is low and that the validation method is suitable for assessing whether GAMLSS models are unstable.

## 1. INTRODUCTION

Normative modeling of measurable features enables both 1.) chartings of population growth and development while also 2.) providing a large comparative reference group to benchmark other individuals and groups.^1^ Recently, this concept has expanded into neuroimaging by modeling derived quantitative measures of brain structure across the lifespan to create human brain charts using magnetic resonance imaging (MRI).^2^ While initially only charts for global brain features were created, more recent work has extended to measures such as cortical thickness^3^ and surface area^4^. These brain charts have been shown to be increasingly useful in detecting anomalies in individuals as well as identifying differences across clinical cohorts.^2,5^

Although the majority of these brain charts focus on gray matter (GM) and whole brain measurements, white matter (WM) remains an important part of brain anatomy, with alterations in WM integrity being linked to function, disease, and normal aging.^6–8^ Recent work from our group has extended brain charts into white matter (WM) microstructure and macrostructure using diffusion-weighted MRI (dMRI) in order to provide a population reference analogous to the whole brain and GM brain charts.^9^ These WM brain charts were created using generalized additive models for location, scale, and shape (GAMLSS).^10^ GAMLSS is a flexible framework that can appropriately model the more complex distributions of quantitative WM measures^11^, as it is supports a variety of distributional families and has the capability to model the mean, spread, skew, and kurtosis of distributions. Further, GAMLSS has been identified by the World Health Organization (WHO) as an appropriate model for creating normative growth charts.^1^

While these WM brain charts have demonstrated their utility in modeling trajectories of WM and identifying abnormalities in clinical cohorts, analytic assessment of model parameter variability at specific timepoints in the lifespan is not commonly examined in other brain charts. Such an approach would be beneficial for assessing model variability, as analytic methods are more computationally efficient, especially given the scale of data being used for normative brain charts.^2,9^ We posit that a validation of the model stability and confidence will provide insight into the reliability of GAMLSS analysis of these normative features. To that end, we conduct an analytic assessment of the variance in these WM brain chart models. Using the models and data, we compare this analytic assessment to an empirical assessment of model stability using a bootstrapping procedure. This comparison between two different methods of variability estimates, analytic and empirical, allows us to determine if the analytic method is an appropriate way to assess model stability of WM brain charts using the GAMLSS framework: analytical approaches for model uncertainty assessment are directly grounded in model assumptions. On the other hand, empirical estimates do not rely on analytic formulas; they are reliable regardless of model mis-specification or violation of assumptions, unlike analytic approaches.

## 2. METHODS

### 2.1 Data

We use the data from our previous work^9^, consisting of *N* = 21,699 typically developing and aging participants from 42 different datasets. (Figure 1) GAMLSS models are fit on microstructural measures of fractional anisotropy (FA), mean diffusivity (MD), axial diffusivity (AD), and radial diffusivity (RD) and macrostructural measures of volume, surface area, and average length (seven total features) for the 72 WM tracts defined by TractSeg^12^, resulting in 504 different WM brain chart models. In the interest of brevity, details of these datasets, preprocessing steps, and quality control processes have been previously described.^9,13,14^

**Figure 1.**
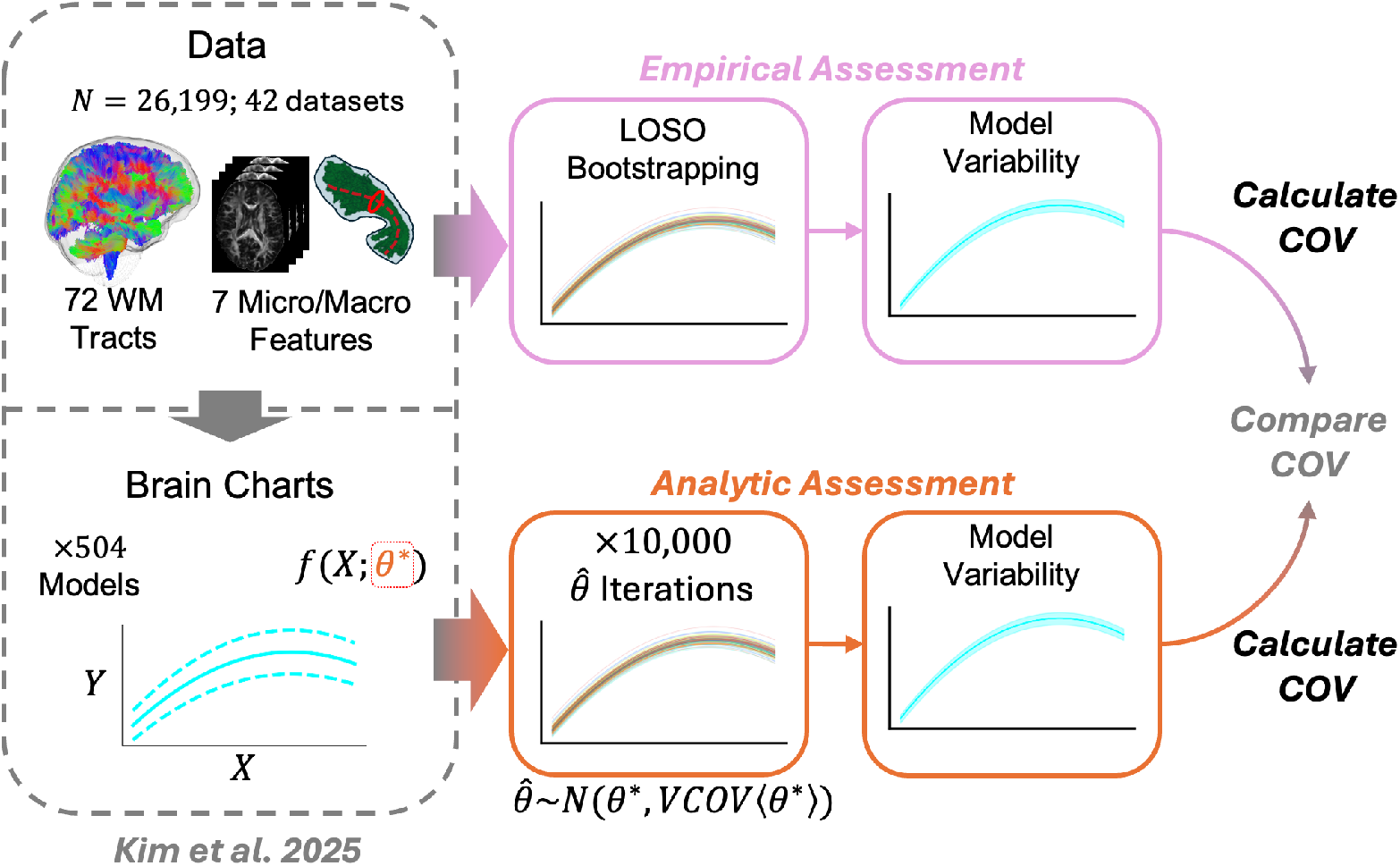
We take the data and WM brain chart models from our previous work to assess model variability. Specifically, we implement an analytic calculation for the variance-covariance matrix of the model parameters *θ*, then use this matrix along with the estimated values for *θ*, or 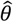, to bootstrap 10,000 different iterations, 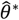, for each model. We also perform leave-one-study-out (LOSO) bootstrapping for each model. For both methods we calculate the coefficient of variation (COV) across the lifespan for each model to compare variabilities.

### 2.2 Variance Estimates

#### The Analytical Approach

For the GAMLSS framework, we estimate the terms, *μ, σ, ν, τ* that parameterize a given distributional family *D*(*μ, σ, ν, τ*) as functions of the independent variables *X*, where the dependent variable *Y* is distributed according to *D*(*μ, σ, ν, τ*).^10^ This framework offers a high degree of flexibility, allowing modeling of random and non-linear effects of our independent variables. For the GAMLSS specification used in our prior work^9^, *D* follows the generalized gamma distribution, with the probability density function *GG*(*x*; *μ, σ, ν*),

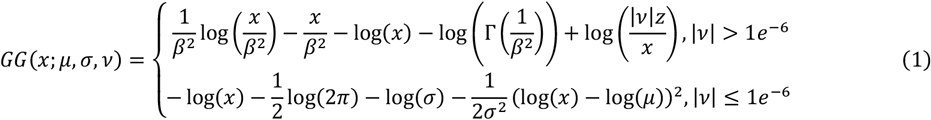

Where

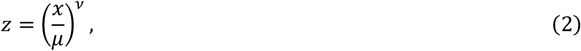

And

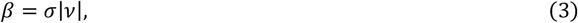

which is a re-parameterization used in the GAMLSS framework.^10^ The generalized gamma distributional parameters, *μ, σ*, and *ν*, are modeled as:

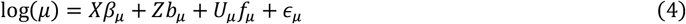

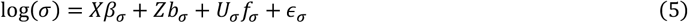

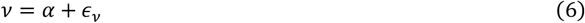

where *β*_*μ*_, *β*_*σ*_ are the vectors of fixed parametric terms for intercept and sex for log(*μ*) and log(*σ*) respectively; *X* is the design matrix for *β*_*μ*_, *β*_*σ*_; *b*_*μ*_, *b*_*σ*_ are the vectors of random intercepts corresponding to dataset for log(*μ*) and log(*σ*) respectively; *Z* is the design matrix for *b*_*μ*_, *b*_*σ*_; *f*_*μ*_, *f*_*σ*_ are the vectors of fractional polynomial coefficients describing non-linear age effects for log(*μ*) and log(*σ*) respectively; *U*_*μ*_, *U*_*σ*_ are the design matrices for *f*_*μ*_, *f*_*σ*_ respectively; *α* is the intercept term for *ν*; and *ϵ*_*μ*_, *ϵ*_*σ*_, *ϵ*_*ν*_ correspond to the error for log(*μ*), log(*σ*), and *ν* respectively. Even though the GAMLSS brain charts are estimated with multiple datasets, each with its own random effect values, the overall trajectories are computed setting the random effect predictions to zero. Thus, to estimate the variance of the lifespan trajectories we need only estimate the joint variance-covariance matrix *V* of the fixed and fractional polynomial estimators 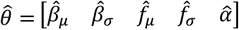.

The GAMLSS fitting algorithm is based on backfitting^10^, where each vector of parameters in 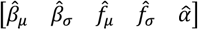 is estimated independently. As a result, each parameter vector has its own separate weight matrix associated with the fit for those parameters only, prohibiting a direct calculation of *V*. However, we can still estimate *V* as the inverse of the Hessian matrix *H* for the negative log likelihood *l* of the GAMLSS model:

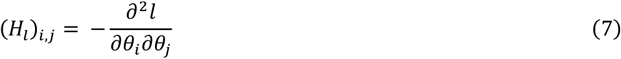

where (*H*_*l*_)^−1^_*i*,*j*_ represents the variance of the *i*^*th*^ parameter when *i* = *j* and the covariance between the *i*^*th*^ and the *j*^*th*^ parameter when *i* ≠ *j*. For numerical stability, we calculate the pseudoinverse of *H*_*l*_. We model each parameter 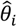 as a random variable that is distributed normally around the true value of the parameter *θ_i_*. If we have *k* parameters in 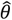, then the *k*-dimensional vector 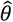 is distributed normally with (*H*_*l*_)^−1^ = *V* as the variance structure of the distribution.

#### The Empirical Approach

For an estimate of the empirical variance, we perform leave-one-study-out (LOSO) bootstrapping as performed in Bethlehem et al.^2^ for each of our 504 models. As we have 42 datasets, we refit 21,168 different bootstrapped models for our empirical estimates. Model fitting was performed using code in the Docker for WM brain charts.^15^ All other experiments were performed using Python 3.9, using the *jax* v0.4.30 library^16^ for calculation of *H*_*l*_, *scipy* v1.13.1^17^ for all statistical tests, and *matplotlib* v3.9.2^18^ and *seaborn* v0.13.2^19^ for quantitative plotting. Tractography plots were made with Python 3.10 using *open-dive*.^20^

### 2.3 Compare Model Variabilities

To estimate the analytic variability of our fit model estimates 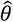, we sample 10,000 times from the *k*-dimensional Gaussian and compute the coefficient of variation (COV) for each model *M*_*s*_ as:

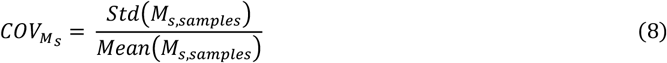

For empirical variability estimates, we calculate COV across all LOSO bootstraps. As model variability can change depending on the age, we compute the COV at 1,000 different linearly spaced points across the lifespan from 0.2 to 90 years of age to assess variability across a range of the age covariate. We then check agreement of the COV averaged across the age range between the analytic and empirical estimates to determine if the analytic method is appropriate for modeling the variability of the GAMLSS WM brain charts.

## 3. RESULTS

For all 504 models, consisting of 72 WM tracts and 7 features, we calculate the model COV at 1,000 linearly spaced points across the lifespan from 0.2 to 90 years of age using 10,000 different samples of 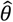 for each model. We observe that across all models, the analytic COV is lower than 5% for the majority of the lifespan (Figure 2, top), with only 2 out of 504 models having larger than 5% COV at the beginning of the lifespan (younger than 0.25 years). Empirical estimates of variance show a similar pattern of low COV across the majority of the lifespan, with 7 out of 504 models having larger than 5% COV during infancy (Figure 2, middle).

**Figure 2.**
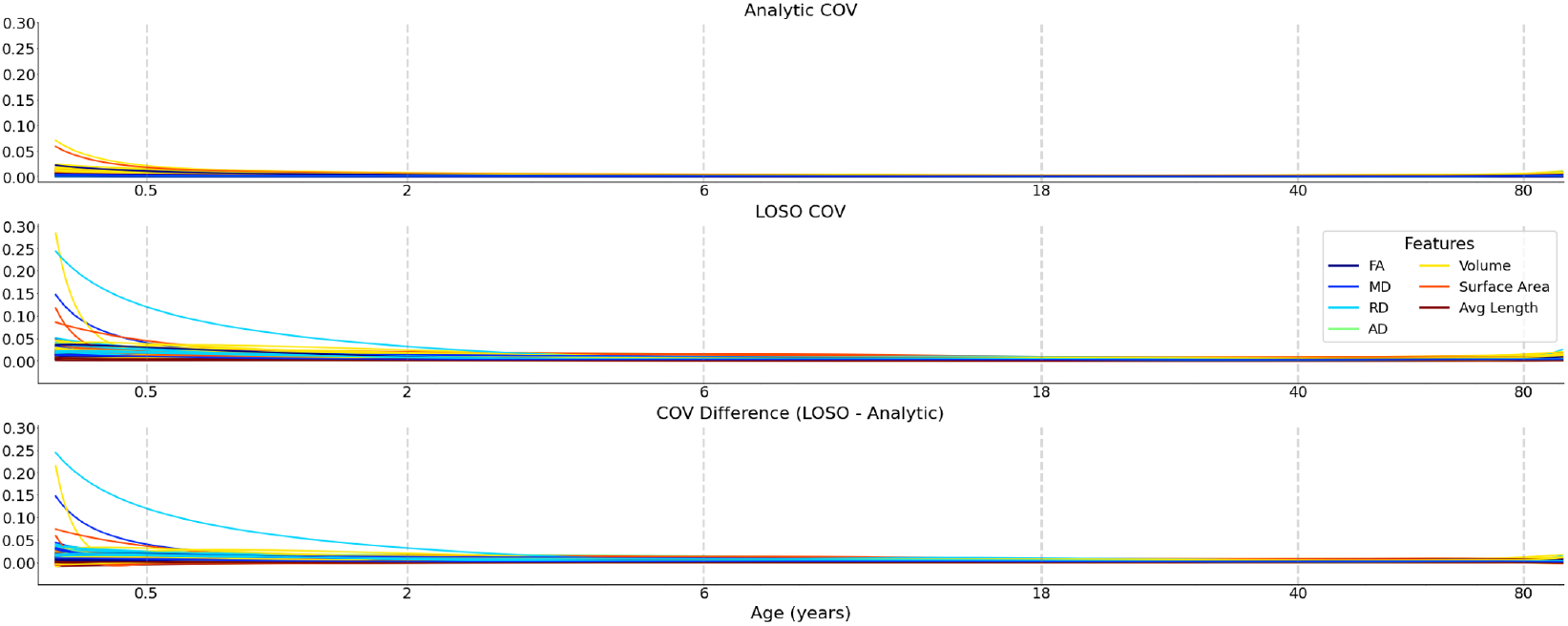
(Top) Analytic variability is relatively low (<5%) for all models across the majority of the lifespan. The COV is highest during infancy, sharply decreasing thereafter. (Middle) Empirical variability is also low across the majority of the lifespan, with some models having relatively high variability during infancy. (Bottom) Most models have strong agreement between both methods across the lifespan, with a few models having higher COV during infancy. Age is plotted with log-spacing to highlight the rapid changes at the beginning of the lifespan.

To directly compare agreement between the analytic and empirical covariance estimates, we average the COV for analytic and empirical estimates of each model across the lifespan. Both methods of estimating the variance have larger averaged COV values for macrostructural features than microstructural features for most tracts (Figure 3). For the empirical analysis with LOSO bootstrapping, macrostructural features have the largest averaged COV for 63 of 72 tracts, whereas the analytic assessment shows that macrostructural features achieve the largest average COV in 71 of 72 tracts. Specifically, tract volume, followed by surface area, is the feature with the largest averaged COV for both methods (Figure 4, left) On the other hand, in the analytic assessment, AD exhibits the smallest average COV for 47 tracts, followed by MD for 13 tracts. Conversely, in the empirical assessment, average length exhibits the smallest average COV for 48 tracts, followed by AD for 15 tracts. (Figure 4, right)

**Figure 3.**
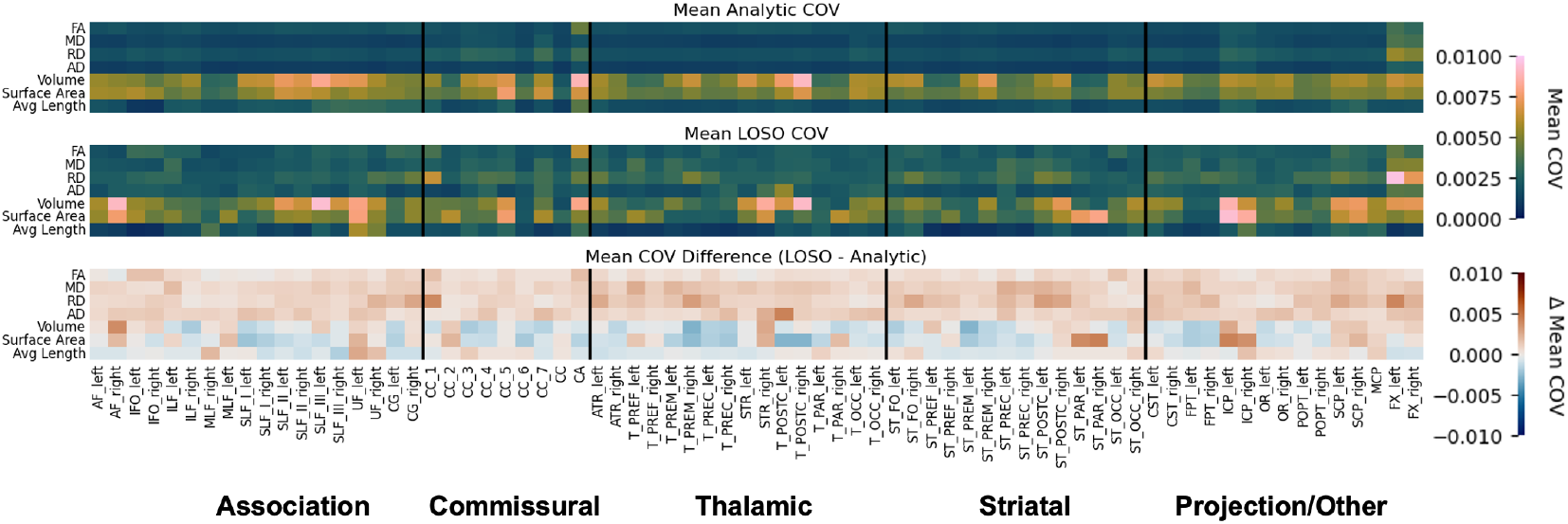
(Top) The average analytic COV across all ages is typically higher for macrostructural measures of volume and surface area than other measures. Tracts that are thinner or smaller, like the fornix (FX_left, FX_right) or anterior commissure (CA), tend to have higher variability than other tracts. (Middle) Empirical COV follows a similar pattern. (Bottom) When compared to empirical COV, analytic COV is underestimated for microstructural measures, but often overestimated for macrostructural measures.

**Figure 4.**
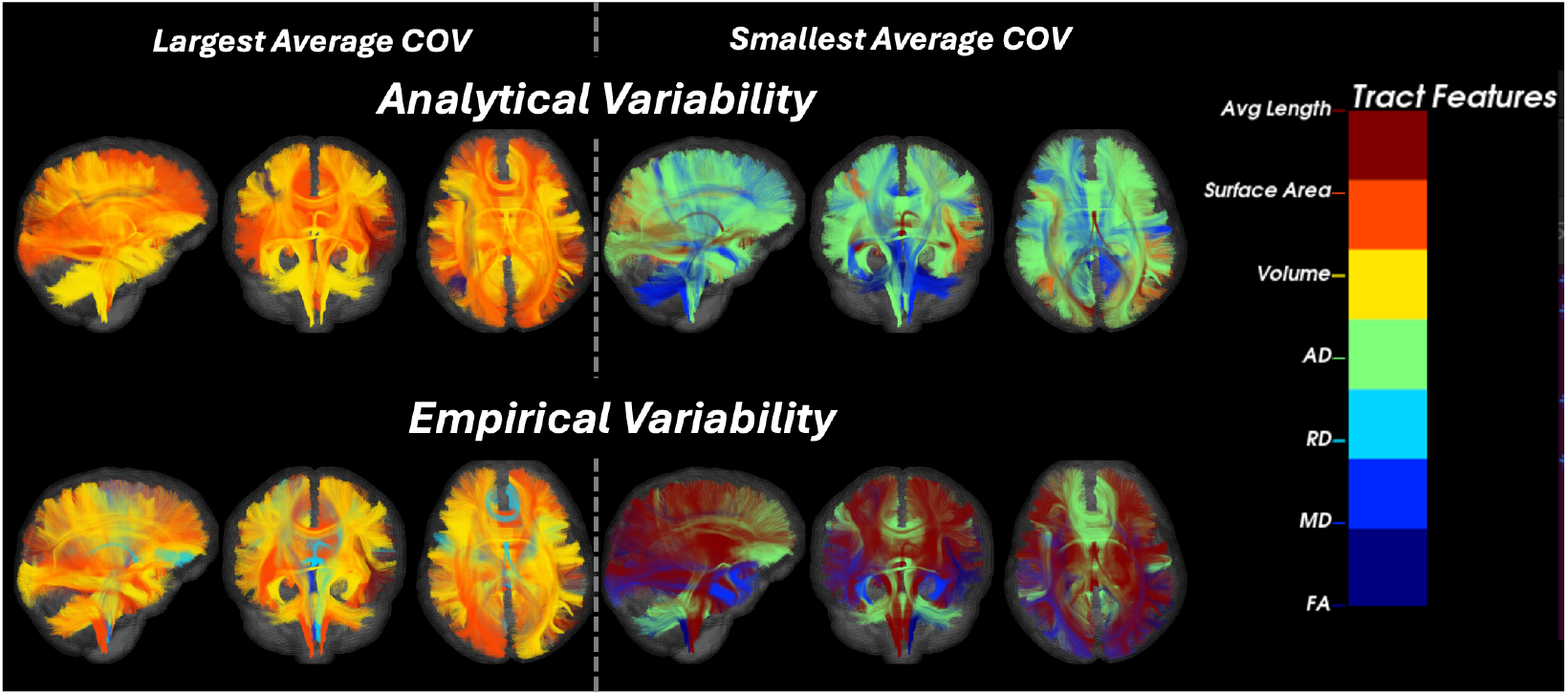
(Left) Across the lifespan, the feature exhibiting the largest analytic and empirical COV is predominantly macrostructural for nearly all tracts. This is true of both the analytic and empirical methods, with this agreement suggesting that the analytic method is appropriate to assess model variability. Volume commonly emerges as the feature with the highest cumulative COV, followed by surface area. (Right) Conversely, the feature with the smallest analytic COV is typically microstructural, with most tracts having AD as the model with the smallest COV, followed by MD. However, for empirical COV, average length is frequently the feature with the smallest COV, followed by AD. Tracts are colored by the respective features with the largest and smallest average COV.

A Bland-Altman plot indicates strong agreement between the analytic and empirical estimates of variability, with 94.64% of data points falling between ±1.96 standard deviations of the mean difference (Figure 5A). Additionally, a Pearson correlation between the two methods showed an agreement coefficient of 0.776, suggesting moderate to high agreement (Figure 5B).

**Figure 5.**
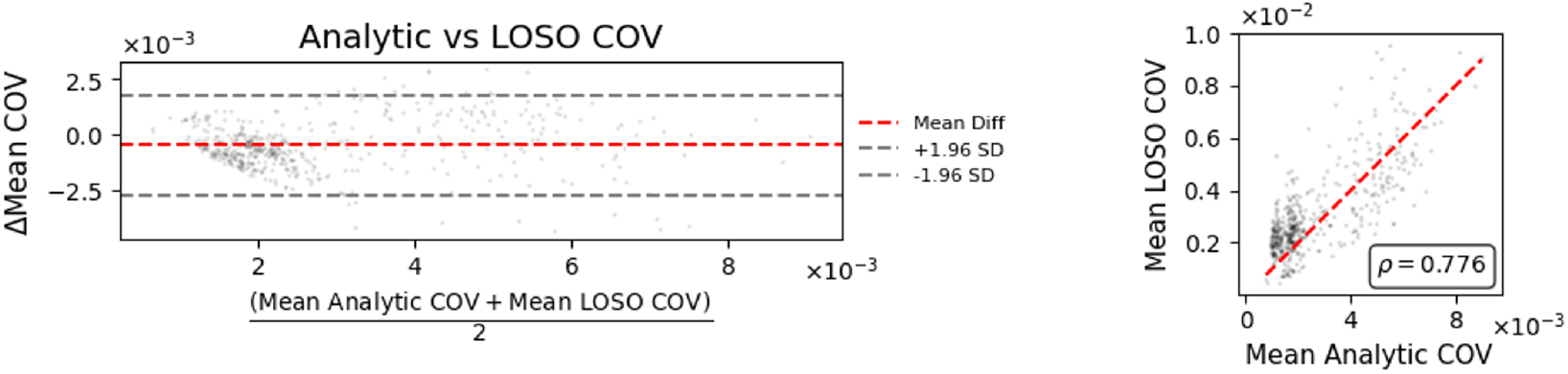
(Left) The analytic and bootstrap methods show moderate to large agreement, suggesting that the GAMLSS model variability is stable and well-approximated by both asymptotic theory and empirical resampling methods. (Right) A Pearson correlation coefficient of 0.776 suggests moderate agreement. There is a high data density in the lower left of the plot, suggesting that even when there is not perfect agreement between the two methods, the COV is still low for both across most models. Further, for extremely confident models with the lowest COV, the analytic approach under-estimates variability.

## 4. DISCUSSION AND CONCLUSION

Nearly 95% of models fall within ±1.96 standard deviations of the mean difference between the analytic and empirical estimates, in addition to a Pearson correlation coefficient of 0.776 (Fig 5). This consistency in variance estimates suggests that the analytic variability is an appropriate method for assessing model stability. However, when the models are very stable, having low empirical COV, the analytic variance estimates further underapproximate the variability. Conversely, when looking at age-specific COV, the disagreement is especially pronounced at the youngest age ranges (< 1 year) for specific models, where there is a small amount of data relative to other age ranges. At these ages, the model experiences edge effects due to being unbounded on the left age extreme^21^ and has data that are inherently lower in signal to noise ratio (SNR)^22^, both of which could be contributing to the higher empirical variability.

A low COV suggests that the WM brain charts are stable and provide reliable estimates for individual benchmarking and lifespan trajectories. Most models remain low in COV (<5% COV) across the entire age range (Figure 2). We observe higher variability in smaller and thinner tracts like the Fornix (FX_left, FX_right) and anterior commissure (CA), which is consistent with results from other multisite studies.^23^ Further, most identified developmental and aging milestones in WM occur at older ages, whereas the few models with higher COV occurred during infancy (Figure 3). This signifies that the brain charts appropriately estimate trajectories at key points in the lifespan. Future work could investigate if including more data at these infant age ranges will increase the model reliability, or if the higher variability is inherent to the low SNR nature of the data. Additionally, the GAMLSS framework for building brain charts could be compared to deep learning-based methods to determine if purely data-driven methods for normative modeling^24^ estimate similar variability to both the analytic and empirical assessments.

**This work has not been submitted for presentation or publication elsewhere**.

## 6. ACKNOWLEDGEMENTS

This work was supported in part by the National Institute of Health through NIH awards K01-EB032898 (Schilling) and K01-AG073584 (Archer), grant number 1R01EB017230-01A1 (Landman), K24-AG046373 (Jefferson), and ViSE/VICTR VR3029, UL1-TR000445, and UL1-TR002243. This work was supported by NIA grants R01-AG034962 (Jefferson), R01-AG056534 (Jefferson), R01-AG062826, and Alzheimer’s Association IIRG-08-88733 (Jefferson) and the NICHD, R01-HD114489 (Vinci-Booher). This work was supported by the Alzheimer’s Disease Sequencing Project Phenotype Harmonization Consortium (ADSP-PHC) that is funded by NIA (U24 AG074855, U01 AG068057 and R01 AG059716). This work was supported by the Department of Defense through grant number HT94252410563. This work was conducted in part using the resources of the Advanced Computing Center for Research and Education (ACCRE) at Vanderbilt University, Nashville, TN. We appreciate the National Institute of HealthS10 Shared Instrumentation grant 1S10OD020154-01, and grant 1S10OD023680-01 (Vanderbilt’s High-Performance Computer Cluster for Biomedical Research). This work was also supported in part by Intramural Research Program of the National Institute on Aging, NIH.

BABIES and ABC datasets were supported by the Jacobs Foundation Early Career Research Fellowship (2017-1261-05) (Humphreys); National Science Foundation CAREER Award (2042285) (Humphreys); Brain and Behavior Research Foundation John and Polly Sparks Foundation Investigator Award (29593) (Humphreys); Vanderbilt Institute for Clinical and Translational Research Grant (VR53419) (Humphreys); Vanderbilt Strong Grant; Vanderbilt Kennedy Center Grant (Humphreys); National Institute of Mental Health (R01MH129634) (Humphreys).

This work was supported by the NIH NINDS through award numbers R01 NS108445 and R01 NS110130 (Morgan), and R01 NS134625 and R01 NS112252 (Englot). Data collected from the MORGAN dataset were acquired at Vanderbilt University, with the aim of studying cognitive patterns in patients with epilepsy both before and after clinical treatment.

This work was supported by the NIH Eunice Kennedy Shriver National Institute Of Child Health & Human Development through grant number F31 HD104385 (Nguyen), P50HD103537 (PI: Neul), R37 HD095519 MERIT Award, R01 HD089474, R01 HD109151, R01 HD067254, and R01 HD044073 (Cutting). This work was also supported by the NIH NINDS through grant number R01 NS049096 (Cutting).

This research was supported in part by the Intramural Research Program of the National Institutes of Health (NIH). The contributions of the NIH author(s) were made as part of their official duties as NIH federal employees, are in compliance with agency policy requirements, and are considered Works of the United States Government. However, the findings and conclusions presented in this paper are those of the author(s) and do not necessarily reflect the views of the NIH or the U.S. Department of Health and Human Services.

Research reported in this publication was supported by NIGMS of the National Institutes of Health under award number T32GM007347 and T32GM152284.

Data and/or research tools used in the preparation of this manuscript were obtained from the National Institute of Mental Health (NIMH) Data Archive (NDA). NDA is a collaborative informatics system created by the National Institutes of Health to provide a national resource to support and accelerate research in mental health. This manuscript reflects the views of the authors and may not reflect the opinions or views of the NIH or of the Submitters submitting original data to NDA.

The Pediatric Imaging, Neurocognition, and Genetics (PING) dataset was collected and released openly to contribute to the assessment of typical brain development in a pediatric sample (RC2DA029475-01), (https://www.sciencedirect.com/science/article/pii/S1053811915003572).

The data used in this study come from the Human Connectome Project, which aims to map the structural connections and circuits of the brain and their relationships to behavior by acquiring high-quality magnetic resonance images. We used diffusion MRI data from the Baby Connectome Project (HCPBaby), the Human Connectome Project Development (HCPD) study, the Human Connectome Project Young Adult (HCP) study, and the Human Connectome Project Aging (HCPA) study.

Research data from the Infant Brain Imaging Study (IBIS) dataset come from the IBIS Autism project, a collaborative effort by investigators to conduct a longitudinal MRI/DTI and behavioral study of infants at high risk for autism (R01HD055741-01).

The Enhanced NKI-RS is a large cross-sectional sample of brain development, maturation and aging, that is currently funded by the NIMH (BRAINS R01MH094639-01; PI Milham) and Child Mind Institute (PI Milham) to characterize 1000 community-ascertained participants using state-of-the-art multiplex imaging-based resting state fMRI (R-fMRI) and diffusion tensor imaging (DTI), genetics, and a broad neurobehavioral phenotypic characterization protocol. Data were acquired from the DSI studio website: (https://brain.labsolver.org/nki_rockland.html).

The Healthy Brain Network (HBN) is an ongoing initiative focused on building a biobank of data from 10,000 children and adolescents (ages 5-21) in the New York City area (https://www.nature.com/articles/sdata2017181). Data were acquired from the DSI studio website: (https://brain.labsolver.org/hbn.html).

Data collection and sharing for this project was provided by the Cambridge Centre for Ageing and Neuroscience (CamCAN). CamCAN funding was provided by the UK Biotechnology and Biological Sciences Research Council (grant number BB/H008217/1), together with support from the UK Medical Research Council and University of Cambridge, UK. Data used in the preparation of this work were obtained from the CamCAN repository (available at http://www.mrc-cbu.cam.ac.uk/datasets/camcan/). The Cambridge Centre for Ageing and Neuroscience (Cam-CAN) is a large-scale collaborative research project at the University of Cambridge.

VMAP data were collected by Vanderbilt Memory and Alzheimer’s Center Investigators at Vanderbilt University Medical Center. VMAP began in 2012 with the goal of investigating vascular health and brain aging.

BLSA is a prospective cohort study with continuous enrollment that began in 1958. Comprehensive data from BLSA are available upon request by a proposal submission through the cohort website (www.blsa.nih.gov). The BLSA is supported by the Intramural Research Program of the National Institute on Aging, NIH.

The BIOCARD study is designed to identify biomarkers associated with progression from normal cognitive status to cognitive impairment or dementia, with a particular focus on Alzheimer’s Disease.

Data collection and sharing for ADNI were supported by National Institutes of Health Grant U01-AG024904 and Department of Defense (award number W81XWH-12-2-0012). ADNI is also funded by the National Institute on Aging, the National Institute of Biomedical Imaging and Bioengineering, and through generous contributions from the following: AbbVie, Alzheimer’s Association; Alzheimer’s Drug Discovery Foundation; Araclon Biotech; BioClinica, Inc.; Biogen; Bristol-Myers Squibb Company; CereSpir, Inc.; Cogstate; Eisai Inc.; Elan Pharmaceuticals, Inc.; Eli Lilly and Company; EuroImmun; F. Hoffmann-La Roche Ltd and its affiliated company Genentech, Inc.; Fujirebio; GE Healthcare; IXICO Ltd.; Janssen Alzheimer Immunotherapy Research & Development, LLC.; Johnson & Johnson Pharmaceutical Research & Development LLC.; Lumosity; Lundbeck; Merck & Co., Inc.; Meso Scale Diagnostics, LLC.; NeuroRx Research; Neurotrack Technologies; Novartis Pharmaceuticals Corporation; Pfizer Inc.; Piramal Imaging; Servier; Takeda Pharmaceutical Company; and Transition Therapeutics. The Canadian Institutes of Health Research is providing funds to support ADNI clinical sites in Canada. Private sector contributions are facilitated by the Foundation for the National Institutes of Health (www.fnih.org). The grantee organization is the Northern California Institute for Research and Education, and the study is coordinated by the Alzheimer’s Therapeutic Research Institute at the University of Southern California. ADNI data are disseminated by the Laboratory for Neuro Imaging at the University of Southern California. Data used in the preparation of this article were obtained from the Alzheimer’s Disease Neuroimaging Initiative (ADNI) database (adni.loni.usc.edu). The ADNI was launched in 2003 as a public private partnership, led by Principal Investigator Michael W. Weiner, MD. The original goal of ADNI was to test whether serial magnetic resonance imaging (MRI), positron emission tomography (PET), other biological markers, and clinical and neuropsychological assessment can be combined to measure the progression of mild cognitive impairment (MCI) and early Alzheimer’s disease (AD). The current goals include validating biomarkers for clinical trials, improving the generalizability of ADNI data by increasing diversity in the participant cohort, and to provide data concerning the diagnosis and progression of Alzheimer’s disease to the scientific community. For up-to-date information, see adni.loni.usc.edu.

Research reported in this publication was supported by the National Institute on Aging of the National Institutes of Health under Award Numbers R01AG054073 and R01AG058533, R01AG070862, P41EB015922 and U19AG078109. The content is solely the responsibility of the authors and does not necessarily represent the official views of the National Institutes of Health.

The NACC database is funded by NIA/NIH Grant U24 AG072122. NACC data are contributed by the NIA-funded ADRCs: P30 AG062429 (PI James Brewer, MD, PhD), P30 AG066468 (PI Oscar Lopez, MD), P30 AG062421 (PI Bradley Hyman, MD, PhD), P30 AG066509 (PI Thomas Grabowski, MD), P30 AG066514 (PI Mary Sano, PhD), P30 AG066530 (PI Helena Chui, MD), P30 AG066507 (PI Marilyn Albert, PhD), P30 AG066444 (PI John Morris, MD), P30 AG066518 (PI Jeffrey Kaye, MD), P30 AG066512 (PI Thomas Wisniewski, MD), P30 AG066462 (PI Scott Small, MD), P30 AG072979 (PI David Wolk, MD), P30 AG072972 (PI Charles DeCarli, MD), P30 AG072976 (PI Andrew Saykin, PsyD), P30 AG072975 (PI David Bennett, MD), P30 AG072978 (PI Ann McKee, MD), P30 AG072977 (PI Robert Vassar, PhD), P30 AG066519 (PI Frank LaFerla, PhD), P30 AG062677 (PI Ronald Petersen, MD, PhD), P30 AG079280 (PI Eric Reiman, MD), P30 AG062422 (PI Gil Rabinovici, MD), P30 AG066511 (PI Allan Levey, MD, PhD), P30 AG072946 (PI Linda Van Eldik, PhD), P30 AG062715 (PI Sanjay Asthana, MD, FRCP), P30 AG072973 (PI Russell Swerdlow, MD), P30 AG066506 (PI Todd Golde, MD, PhD), P30 AG066508 (PI Stephen Strittmatter, MD, PhD), P30 AG066515 (PI Victor Henderson, MD, MS), P30 AG072947 (PI Suzanne Craft, PhD), P30 AG072931 (PI Henry Paulson, MD, PhD), P30 AG066546 (PI Sudha Seshadri, MD), P20 AG068024 (PI Erik Roberson, MD, PhD), P20 AG068053 (PI Justin Miller, PhD), P20 AG068077 (PI Gary Rosenberg, MD), P20 AG068082 (PI Angela Jefferson, PhD), P30 AG072958 (PI Heather Whitson, MD), P30 AG072959 (PI James Leverenz, MD).

This research has been conducted using the UK Biobank resource, application 16315.

Data contributed from MAP/ROS/MARS was supported by NIA R01AG017917, P30AG10161, P30AG072975, R01AG022018, R01AG056405, UH2NS100599, UH3NS100599, R01AG064233, R01AG15819 and R01AG067482, and the Illinois Department of Public Health (Alzheimer’s Disease Research Fund). Data can be accessed at www.radc.rush.edu. More information about participant demographics and study information can be found here: https://www.rushu.rush.edu/research-rush-university/departmental-research/rush-alzheimers-disease-center/rush-alzheimers-disease-center-research/epidemiologic-research.

The data contributed from the Wisconsin Registry for Alzheimer’s Prevention was supportedby NIA AG021155, AG0271761, AG037639, and AG054047.

We use generative AI to create code segments based on task descriptions, as well as debug, edit, and autocomplete code. Additionally, generative AI technologies have been employed to assist in structuring sentences and performing grammatical checks. It is imperative to highlight that the conceptualization, ideation, and all prompts provided to the AI originate entirely from the authors’ creative and intellectual efforts. We take accountability for the review of all content generated by AI in this work.

Data collection and sharing for this project was provided by the Centre for Attention, Learning and Memory (CALM). CALM funding was provided by the UK Medical Research Council and University of Cambridge, UK. Data used in the preparation of this work were obtained from CALM resource – https://calm.mrc-cbu.cam.ac.uk/. The study protocol is reported in Holmes et al. (2019).

Data used in the preparation of this work were obtained from the International Consortium for Brain Mapping (ICBM) database (www.loni.usc.edu/ICBM). The ICBM project (Principal Investigator John Mazziotta, M.D., University of California, Los Angeles) is supported by the National Institute of Biomedical Imaging and BioEngineering. ICBM is the result of efforts of co-investigators from UCLA, Montreal Neurologic Institute, University of Texas at San Antonio, and the Institute of Medicine, Juelich/Heinrich Heine University -Germany. Data collection and sharing for this project was provided by the International Consortium for Brain Mapping (ICBM; Principal Investigator: John Mazziotta, MD, PhD). ICBM funding was provided by the National Institute of Biomedical Imaging and BioEngineering. ICBM data are disseminated by the Laboratory of Neuro Imaging at the University of Southern California.

The dataset that we refer to as UTAustin in this manuscript is a longitudinal neuroimaging dataset on language processing in children ages 5, 7, and 9 years old collected at The University of Texas at Austin. The dataset is openly available from Openneuro here: https://openneuro.org/datasets/ds003604/versions/1.0.7. We use version 1.0.7.

The Queensland Twin Adolescent Brain (QTAB) Project was established with the purpose of promoting the conduct of health-related research in adolescence. The QTAB dataset comprises multimodal neuroimaging, as well as cognitive and mental health data collected in adolescent twins over two sessions. The MRI protocol consisted of T1-weighted (MP2RAGE), T2-weighted, FLAIR, high-resolution TSE, SWI, resting-state fMRI, DWI, and ASL scans. The QTAB project resource was produced as a result of i) the goodwill and contribution of 422 twin/triplet participants and their parents, ii) funding from the National Health and Medical Research Council, Australia (APP1078756) and the Queensland Brain Institute, University of Queensland, iii) access to several key resources, including the Centre for Advanced Imaging, the Human Studies Unit, Institute of Molecular Bioscience, and the Queensland Cyber Infrastructure Foundation, at the University of Queensland, local and national twin registries at the QIMR Berghofer Medical Research Institute and Twin Research Australia, as well as the many assessments made available by researchers worldwide, and iv) was established with the purpose of promoting the conduct of health-related research in adolescence. The imaging data and basic demographics are openly accessibly on Openneuro here: https://openneuro.org/datasets/ds004146/versions/1.0.4. We use version 1.0.4.

The Social Reward and Nonsocial Reward Processing Across the Adult Lifespan: An Interim Multi-echo fMRI and Diffusion Dataset (referred to as TempleSocial in this manuscript) comes from a study that aims to investigate whether older adults have a blunted response to some features of social reward. The dataset is openly accessibly on Openneuro here: https://openneuro.org/datasets/ds005123/versions/1.1.3. In particular, we use version 1.1.3.

The Longitudinal Brain Correlates of Multisensory Lexical Processing in Children study (shortened to Lexical in this manuscript) aims to explore developmentally dependent changes in lexical processing for adolescents. The dataset is openly accessibly on Openneuro here: https://openneuro.org/datasets/ds001894/versions/1.4.2. We use version 1.4.2.

The UCLA Consortium for Neuropsychiatric Phenomics LA5c Study (UCLA) is focused on understanding the dimensional structure of memory and cognitive control (response inhibition) functions in both healthy individuals and individuals with neuropsychiatric disorders including schizophrenia, bipolar disorder, and attention deficit/hyperactivity disorder. Neuroimaging data were downloaded from Openneuro here: https://openneuro.org/datasets/ds000030/versions/1.0.0. We use version 1.0.0.

The dataset we refer to as UPennRisk comes from a study at University of Pennsylvania that investigated whether training executive cognitive function could influence choice behavior and brain responses. Neuroimaging data were downloaded from Openneuro here: https://openneuro.org/datasets/ds002843/versions/1.0.1. We use version 1.0.1.

The Dallas Lifespan Brain Study (DLBS) is a longitudinal multi-modal neuroimaging study of the aging mind, which was initiated in 2008 (referred to as Wave 1). Participants returned for two additional waves of data collection with an approximate interval of 4-5 years between waves. The DLBS protocol encompasses various imaging modalities, including structural MRI, diffusion MRI, and functional MRI, as well as comprehensive cognitive and psychosocial assessments. DLBS data can be downloaded from Openneuro here: https://openneuro.org/datasets/ds004856. Specifically, we use version 1.2.0.

The Multisite, Multiscanner, and Multisubject Acquisitions for Studying Variability in Diffusion Weighted Magnetic Resonance Imaging (MASiVar) dataset consists of 319 diffusion scans acquired at 3T from b = 1000 to 3000 s/mm2 across 14 healthy adults, 83 healthy children (5 to 8 years), three sites, and four scanners curated to promote investigation of diffusion MRI variability. In particular, we used only the data coming from healthy children (Cohort IV) for version 2.0.2 of the dataset. Data are available to download from Openneuro here: https://openneuro.org/datasets/ds003416/versions/2.0.2.

Data coming from the Southwestern University (SWU) dataset, part of the Consortium for Reliability and Reproducibility (CoRR), were downloaded via NITRC-IR from the 1000 Functional Connectomes Project. Specifically, the data come from the Emotion and Creativity One Year Retest Dataset subset, comprised of 235 subjects, all of whom were college students. Each subject underwent two sessions of anatomical, resting state fMRI, and DTI scans, spaced one year apart. In order to access the CoRR datasets through NITRC, users must be logged into NITRC at the time of download and registered with the 1000 Functional Connectomes Project / INDI website. More information about this subset can be found here: https://fcon_1000.projects.nitrc.org/indi/CoRR/html/swu_4.html.

The Preschool MRI study in The Developmental Neuroimaging Lab at the University of Calgary (https://www.developmentalneuroimaginglab.ca) uses different magnetic resonance imaging (MRI) techniques to study brain structure and function in early childhood. The study aims to characterize brain development in early childhood, and to offer baseline data that can be used to understand cognitive and behavioural development, as well as to identify deviations from normal development in children with various diseases, disorders, or brain injuries. The MRI techniques used include diffusion tensor imaging (DTI), anatomical imaging, arterial spin labeling (ASL), and resting state functional MRI (rsfMRI). Data can be downloaded from here: https://osf.io/axz5r/.

The Amsterdam Open MRI Collection (AOMIC) is a collection of three datasets with multimodal (3T) MRI data including structural (T1-weighted), diffusion-weighted, and (resting-state and task-based) functional BOLD MRI data, as well as detailed demographics and psychometric variables from a large set of healthy participants. All raw data is publicly available from the Openneuro data sharing platform: ID1000: https://openneuro.org/datasets/ds003097, PIOP1: https://openneuro.org/datasets/ds002785, PIOP2: https://openneuro.org/datasets/ds002790. We use version 1.2.1 for ID1000 and 2.0.0 for PIOP1 and PIOP2.

The Boston Adolescent Neuroimaging of Depression and Anxiety (BANDA) is a study of 215 adolescents ages 14-17, 152 of whom had a current diagnosis of a DSM-5 (APA, 2013) anxious and/or depressive disorder. The BANDA study collected a rich dataset of brain, clinical, and cognitive/neuropsychological measures from these adolescent subjects. The dataset is available to download upon request on the NDA.

We thank Knight ADRC for providing neuroimaging data to us (WASHU dataset). The data contributed through WASHU (Knight ADRC) was supported by grant numbers P30 AG066444, P01 AG03991, and P01 AG026276. For WASHU, Clinical Dementia Ratings (CDRs) are obtained from assessments by experienced clinicians trained in the use of the CDR.

The NACC database is funded by NIA/NIH Grant U24 AG072122. SCAN is a multi-institutional project that was funded as a U24 grant (AG067418) by the National Institute on Aging in May 2020. Data collected by SCAN and shared by NACC are contributed by the NIA-funded ADRCs as follows:

Arizona Alzheimer’s Center - P30 AG072980 (PI: Eric Reiman, MD); R01 AG069453 (PI: Eric Reiman (contact), MD); P30 AG019610 (PI: Eric Reiman, MD); and the State of Arizona which provided additional funding supporting our center; Boston University - P30 AG013846 (PI Neil Kowall MD); Cleveland ADRC - P30 AG062428 (James Leverenz, MD); Cleveland Clinic, Las Vegas – P20AG068053; Columbia - P50 AG008702 (PI Scott Small MD); Duke/UNC ADRC – P30 AG072958; Emory University - P30AG066511 (PI Levey Allan, MD, PhD); Indiana University - R01 AG19771 (PI Andrew Saykin, PsyD); P30 AG10133 (PI Andrew Saykin, PsyD); P30 AG072976 (PI Andrew Saykin, PsyD); R01 AG061788 (PI Shannon Risacher, PhD); R01 AG053993 (PI Yu-Chien Wu, MD, PhD); U01 AG057195 (PI Liana Apostolova, MD); U19 AG063911 (PI Bradley Boeve, MD); and the Indiana University Department of Radiology and Imaging Sciences; Johns Hopkins - P30 AG066507 (PI Marilyn Albert, Phd.); Mayo Clinic - P50 AG016574 (PI Ronald Petersen MD PhD); Mount Sinai - P30 AG066514 (PI Mary Sano, PhD); R01 AG054110 (PI Trey Hedden, PhD); R01 AG053509 (PI Trey Hedden, PhD); New York University - P30AG066512-01S2 (PI Thomas Wisniewski, MD); R01AG056031 (PI Ricardo Osorio, MD); R01AG056531 (PIs Ricardo Osorio, MD; Girardin Jean-Louis, PhD); Northwestern University - P30 AG013854 (PI Robert Vassar PhD); R01 AG045571 (PI Emily Rogalski, PhD); R56 AG045571, (PI Emily Rogalski, PhD); R01 AG067781, (PI Emily Rogalski, PhD); U19 AG073153, (PI Emily Rogalski, PhD); R01 DC008552, (M.-Marsel Mesulam, MD); R01 AG077444, (PIs M.-Marsel Mesulam, MD, Emily Rogalski, PhD); R01 NS075075 (PI Emily Rogalski, PhD); R01 AG056258 (PI Emily Rogalski, PhD); Oregon Health and Science University - P30 AG008017 (PI Jeffrey Kaye MD); R56 AG074321 (PI Jeffrey Kaye, MD); Rush University - P30 AG010161 (PI David Bennett MD); Stanford – P30AG066515; P50 AG047366 (PI Victor Henderson MD MS); University of Alabama, Birmingham – P20; University of California, Davis - P30 AG10129 (PI Charles DeCarli, MD); P30 AG072972 (PI Charles DeCarli, MD); University of California, Irvine - P50 AG016573 (PI Frank LaFerla PhD); University of California, San Diego - P30AG062429 (PI James Brewer, MD, PhD); University of California, San Francisco - P30 AG062422 (Rabinovici, Gil D., MD); University of Kansas - P30 AG035982 (Russell Swerdlow, MD); University of Kentucky - P30 AG028283-15S1 (PIs Linda Van Eldik, PhD and Brian Gold, PhD); University of Michigan ADRC - P30AG053760 (PI Henry Paulson, MD, PhD) P30AG072931 (PI Henry Paulson, MD, PhD) Cure Alzheimer’s Fund 200775 - (PI Henry Paulson, MD, PhD) U19 NS120384 (PI Charles DeCarli, MD, University of Michigan Site PI Henry Paulson, MD, PhD) R01 AG068338 (MPI Bruno Giordani, PhD, Carol Persad, PhD, Yi Murphey, PhD) S10OD026738- 01 (PI Douglas Noll, PhD) R01 AG058724 (PI Benjamin Hampstead, PhD) R35 AG072262 (PI Benjamin Hampstead, PhD) W81XWH2110743 (PI Benjamin Hampstead, PhD) R01 AG073235 (PI Nancy Chiaravalloti, University of Michigan Site PI Benjamin Hampstead, PhD) 1I01RX001534 (PI Benjamin Hampstead, PhD) IRX001381 (PI Benjamin Hampstead, PhD); University of New Mexico - P20 AG068077 (Gary Rosenberg, MD); University of Pennsylvania - State of PA project 2019NF4100087335 (PI David Wolk, MD); Rooney Family Research Fund (PI David Wolk, MD); R01 AG055005 (PI David Wolk, MD); University of Pittsburgh - P50 AG005133 (PI Oscar Lopez MD); University of Southern California - P50 AG005142 (PI Helena Chui MD); University of Washington - P50 AG005136 (PI Thomas Grabowski MD); University of Wisconsin - P50 AG033514 (PI Sanjay Asthana MD FRCP); Vanderbilt University – P20 AG068082; Wake Forest - P30AG072947 (PI Suzanne Craft, PhD); Washington University, St. Louis - P01 AG03991 (PI John Morris MD); P01 AG026276 (PI John Morris MD); P20 MH071616 (PI Dan Marcus); P30 AG066444 (PI John Morris MD); P30 NS098577 (PI Dan Marcus); R01 AG021910 (PI Randy Buckner); R01 AG043434 (PI Catherine Roe); R01 EB009352 (PI Dan Marcus); UL1 TR000448 (PI Brad Evanoff); U24 RR021382 (PI Bruce Rosen); Avid Radiopharmaceuticals / Eli Lilly; Yale - P50 AG047270 (PI Stephen Strittmatter MD PhD); R01AG052560 (MPI: Christopher van Dyck, MD; Richard Carson, PhD); R01AG062276 (PI: Christopher van Dyck, MD); 1Florida - P30AG066506-03 (PI Glenn Smith, PhD); P50 AG047266 (PI Todd Golde MD PhD)

During the preparation of this work, the authors used GitHub Copilot and ChatGPT to assist with code generation, debugging, editing, and sentence restructuring. All AI-generated content was reviewed and edited by the authors, who take full responsibility for the final content of the publication.

